# Towards environmental detection, quantification, and molecular characterization of *Anopheles stephensi* and *Aedes aegypti* from larval breeding sites

**DOI:** 10.1101/2022.09.29.510135

**Authors:** Mojca Kristan, Holly Acford-Palmer, Monica Oliveira Campos, Emma Collins, Jody Phelan, Natalie M. Portwood, Bethanie Pelloquin, Sian Clarke, Jo Lines, Taane G. Clark, Thomas Walker, Susana Campino, Louisa A. Messenger

## Abstract

The invasion and establishment of *An. stephensi* mosquitoes in the Horn of Africa represents a significant regional threat, which may jeopardise malaria control, particularly in urban areas which were formally free from disease transmission. Novel vector surveillance methods are urgently needed, both agnostic to mosquito larval morphology, and simple to implement at the sampling stage. Using new multiplex TaqMan assays, specifically targeting *An. stephensi* and *Ae. aegypti*, we validated the use of environmental DNA (eDNA) for simultaneous vector detection in shared artificial breeding sites. Study findings demonstrated that *An. stephensi* and *Ae. aegypti* eDNA deposited by as few as one second instar larva in 1L of water was detectable. Characterization of molecular insecticide resistance mechanisms, using novel amplicon-sequencing panels for both vector species, was possible from eDNA shed by as few as 32 second instar larvae in 50ml of water. *An. stephensi* eDNA, derived from emergent pupae for 24 hours, was remarkably stable, and still detectable ~2 weeks later. eDNA surveillance has the potential to be implemented in local endemic communities and points of country entry, to monitor the spread of invasive vector species. Further studies are required to validate the feasibility of this technique under field conditions.

## Introduction

Vector-borne diseases (VBDs) account for 17% of all infectious diseases worldwide and are responsible for more than 700,000 deaths annually^1^. In sub-Saharan Africa, malaria, transmitted by *Anopheles* mosquitoes, remains the leading cause of morbidity and mortality (241 million new cases and 627,000 deaths in 2020); despite the widespread deployment of effective vector control measures, especially the use of insecticide-treated nets (ITNs) and indoor residual spraying (IRS), which have averted more than 1.5 billion cases and 7.6 million malaria-related deaths since 2000^2^. Concurrently, approximately 51 million individuals are infected with lymphatic filariasis^3^, a severe cause of disability and stigma, and many arthropod-borne viruses (arboviruses) including, dengue, chikungunya, Rift Valley fever, yellow fever, Zika, o’nyong’nyong and West Nile, circulate unabated among humans, wildlife and livestock across the continent^4,5^. Over 27,000 cases of viruses transmitted by *Aedes* mosquitoes have been reported in West Africa since 2007, but seroprevalence surveys, indicative of prior disease exposure, suggest this is a gross underestimation of the actual arboviral disease burden^6^. While there is growing recognition that parts of Africa are at serious risk for arbovirus outbreaks, the understanding of the current distribution of these diseases is inadequate, as regional VBD surveillance is focused primarily on malaria, detection is usually reactive in response to outbreaks^7^, and robust evidence for efficacious control tools targeting arbovirus vectors is lacking^8^.

Entomological monitoring is an essential part of vector control, providing information on vector species present in an area and supporting data used for risk assessment, planning, implementation, monitoring and the evaluation of control interventions^9^. The World Health Organization (WHO) considers vector surveillance a core component of malaria control programmes^2^. Similarly, a need for arboviral disease control programmes focused on surveillance, vector control and case management has been recognised, especially for dengue^10^. The development of effective disease control strategies requires detailed knowledge of local mosquito vectors. Several different methods can be used for vector surveillance, including larval surveys and adult trapping, and the choice depends on the target vector species, location, purpose of sampling, as well as available resources and infrastructure. However, the expansion of vector species into new geographical areas may be challenging due to markedly different larval habitats and adult behaviour, issues with identifying a new species and potentially varying trapping efficiencies^11,12^.

*Anopheles stephensi* is a highly competent malaria vector whose distribution until 2011 encompassed the Indian subcontinent, parts of South-East Asia and the Arabian Peninsula. Recently it has become an invasive vector species in the horn of Africa, recognised by the WHO as a “biological threat”^13^. *An. stephensi* was first reported in Djibouti in 2012^14^ and since then this species has spread to neighbouring Ethiopia (2016)^15,16^, the Republic of Sudan (2019)^17,18^, Somalia (2019)^19^, Yemen (2021)^19^ and most recently Nigeria (2020)^19^. The invasion of *An. stephensi* in Djibouti was accompanied by an increase in malaria cases from 25 in 2012 to 14,810 in 2017^20^, with outbreaks reported in Djibouti City, including historically malaria-free areas that were considered unsuitable for anopheline breeding sites^14^. Unlike *An. arabiensis*, the main malaria vector species in this region, *An. stephensi* often breeds in urban areas in man-made water containers, buckets, discarded tyres, and water storage tanks for domestic use and construction. These are sites that might not usually be under routine surveillance by the National Malaria Control Programme (NMCP) and if larvae are found, they may be incorrectly identified or simply assumed to be *Aedes* species. Such larval habitats are often used by *Aedes* species, vectors responsible for dengue outbreaks and transmission of other arboviruses, including yellow fever, in Ethiopia^21,22^. *An. stephensi* and *Ae. aegypti* have already been observed in the same larval habitats in eastern Ethiopia^23^. Given the propensity of the former vector species to colonize urban environments quickly, breeding mainly in man-made water containers, mathematical models estimate over 126 million people in cities across Africa are at risk of malaria transmitted by *An. stephensi*^24^ and that annual *Plasmodium falciparum* cases could increase by 50% if no vector control interventions are implemented^25^.

As the spread of *An. stephensi* poses a significant problem to urban populations in Africa, the WHO recommends targeted surveillance to monitor existing populations, dispersal dynamics and to characterize their insecticide resistance profiles to inform prospective vector control strategies^26^. Moreover, dengue has been recognised as a significant public health threat in Ethiopia, where a need for improved vector surveillance and control has been recognised^21^. Novel vector surveillance methods are urgently required, which demand little prior knowledge of mosquito larval morphology and are quick and easy to implement at the sampling stage, with the ability to simultaneously detect the presence of multiple species of interest. Recent advances in molecular techniques are providing new opportunities for surveillance and prevention of VBDs, which can be used to better target vector control strategies. Detection of the traces of genetic material that organisms leave behind in their environment is one possible approach. Environmental DNA (eDNA) found in collected samples may allow us to indirectly confirm the presence of a species of interest and its abundance even when there are no obvious signs of the organism being present^27,28^. eDNA can be extracted from environmental samples, such as water from potential mosquito larval habitats, and can be detected using sensitive molecular techniques such as qPCR. eDNA decay and degradation are environmentally-dependent and due to its nature, detection of eDNA is an indicator of recent presence of the species of interest^29^. This approach can be especially useful for detection of invasive species, including mosquitoes^28,30,31^.

This study assessed the suitability of using eDNA for simultaneous detection of *An. stephensi* and *Ae. aegypti* from the same larval habitats and characterization of their molecular insecticide resistance mechanisms, under controlled laboratory conditions.

## Results

### PCR primer testing and validation

To enable simultaneous detection of *An. stephensi* and *Ae. aegypti* eDNA sampled from shared breeding sites, we designed a multiplex TaqMan assay based on single nucleotide polymorphisms (SNPs) in COX1. Initially the analytical sensitivity, linearity, and dynamic range of the qPCR assay for eDNA detection was estimated using serial dilutions of *An. stephensi* and *Ae. aegypti* gDNA derived from colony strains, tested in technical triplicate. Both qPCR assays produced good linearity (*R*^2^=0.99 for both *An. stephensi* and *Ae. aegypti* assays) and efficiencies of 90.01% and 94.62%, respectively (Figure 1A and B). The LODs (limits of detection) and LOQs (limits of quantification) were determined to be 0.0000154 copies per reaction for individual detection of *An. stephensi* and *Ae. aegypti*. Next, the analytical sensitivity, linearity and dynamic range of the multiplex qPCR assay was evaluated using *An. stephensi* and *Ae. aegypti* gDNA serially diluted in equal proportions. Similarly, the multiplex assay was highly efficient (102.2% and 104.9%for *An. stephensi* and *Ae. aegypti*,respectively) with good linearities (R^2^=0.99 for simultaneous detection of *An. stephensi* and *Ae. aegypti* gDNA). The LoD/LoQs were determined to be 0.0000154 copies/reaction for simultaneous detection of *An. stephensi* and *Ae. aegypti* (Figure 1C).

**Figure 1.**
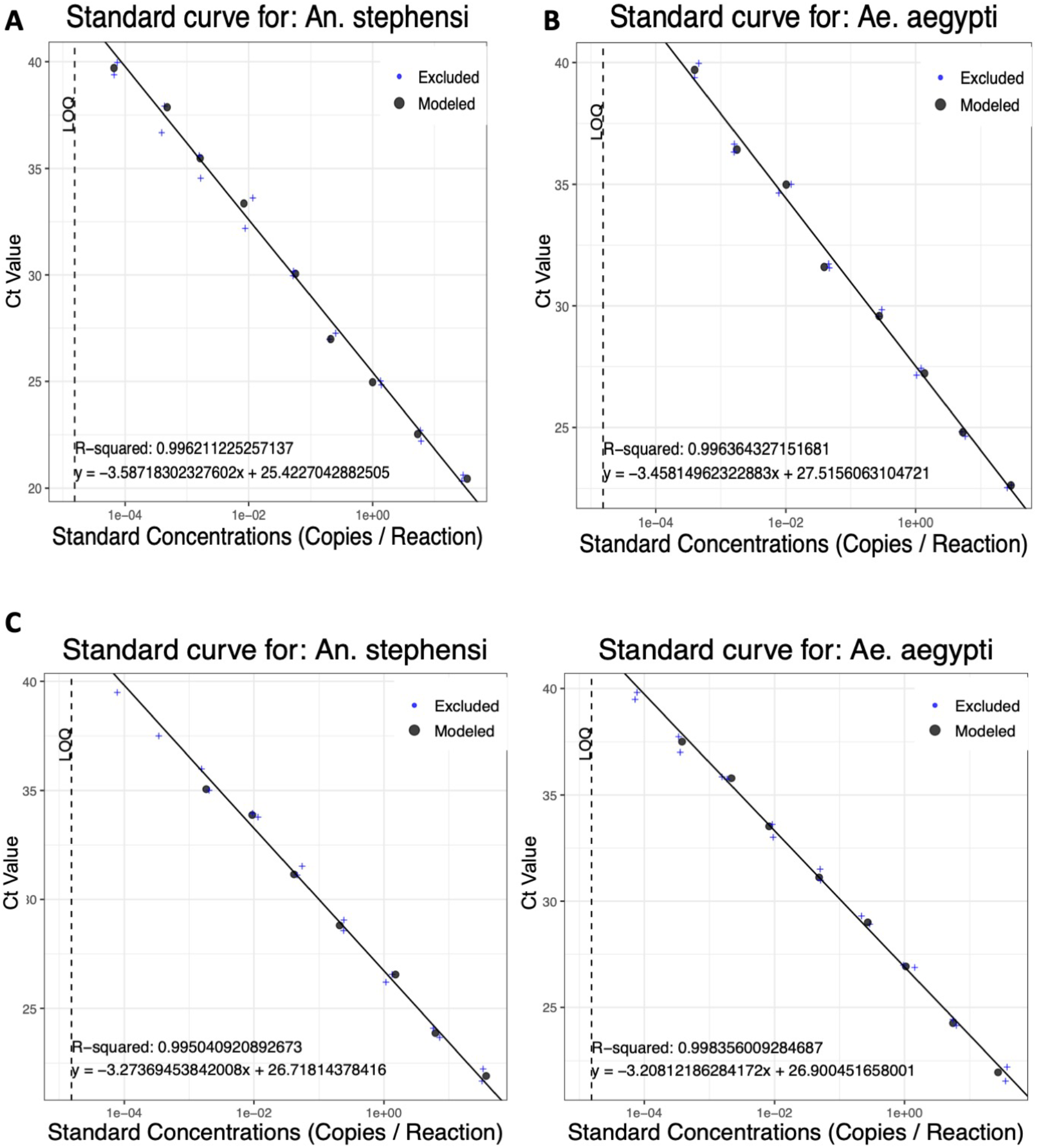
Standard curves across a tenfold dilution series of gDNA for individual detection of *An. stephensi* (A) or *Ae. aegypti* (B) and simultaneous detection of *An. stephensi* and *Ae. aegypti* (C), across a tenfold gDNA dilution series.

The *An. stephensi* and *Ae. aegypti* probes were highly specific to gDNA from each species; no amplification was detected with gDNA from other sympatric or medically important vector species: *An. arabiensis, An. gambiae* s.s., *An. funestus* s.s. or *Culex quinquefasciatus*. All qPCR data are reported in Supplementary File S1.

### Detection of An. stephensi and Ae. aegypti eDNA in artificial breeding sites

To investigate the sensitivity of *An. stephensi* and *Ae. aegypti* eDNA detection, artificial breeding sites of 50ml or 1L of water were simulated with known numbers of second instar larvae (Figure 2).

**Figure 2.**
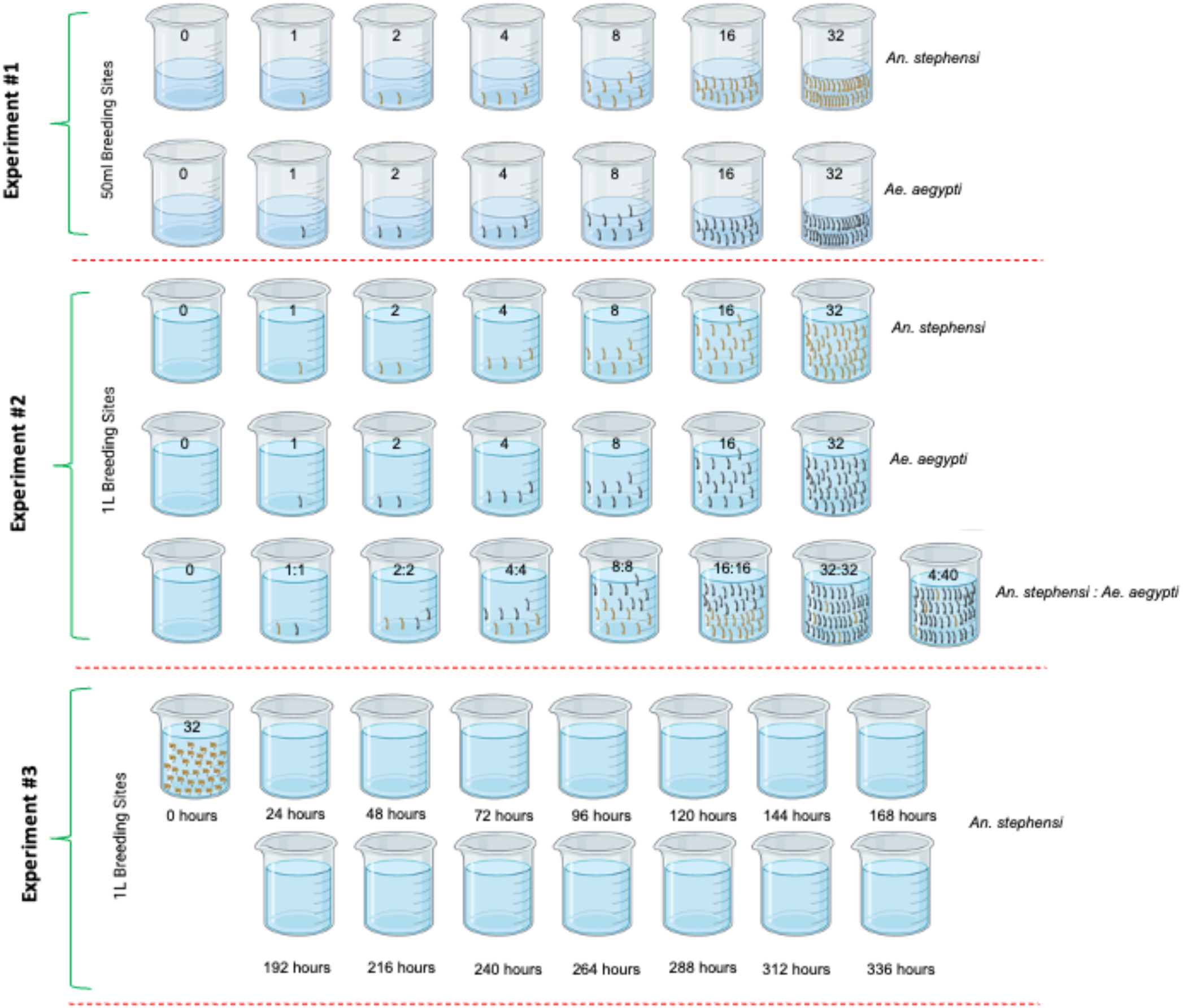
Design of experiments to investigate the impact of different environmental conditions on *An. stephensi* and *Ae. aegypti* eDNA detection. Numbers of second instar larvae are denoted on each artificial breeding site. Experiment 1 investigated eDNA detection of different densities of *An. stephensi* and *Ae. aegypti* in 50ml artificial breeding sites. Experiment 2 investigated eDNA detection of different densities of *An. stephensi* and *Ae. aegypti* in 1L artificial breeding sites, including different ratios of *An. stephensi:Ae. aegypti* co-habiting the same breeding site. Experiment 3 investigated the rate of eDNA degradation. *An. stephensi* pupae were left in each breeding site for 24 hours, prior to manual removal of all emerged adults, remaining pupae, and pupal skins; eDNA detection was performed for the following 14 days. Figure created with BioRender.com.

In 50ml artificial breeding sites (Figure 2, Experiment 1), a dose response was evident for detection levels of both vector species; with increasing densities of larvae, cycle threshold (Ct) values began to decrease significantly (Figure 3). The average Ct values for *An. stephensi* for each larval density were 1 larva: 31.89 [95% confidence interval (CI): 31.35-32.42]; 2 larvae: 27.36 [95% CI: 24.72-29.99]; 4 larvae: 26.74 [95% CI: 25.75-27.73]; 8 larvae: 25.42 [95% CI: 23.77-27.07]; 16 larvae: 21.96 [95% CI: 21.09-22.83]; 32 larvae: 21.01 [95% CI: 20.09-21.92] (Figure 3A). The average Ct values for *Ae. aegypti* for each larval density were 1 larva: 33.66 [95% CI: 31.15-36.16]; 2 larvae: 29.11 [95% CI: 27.19-31.03]; 4 larvae: 29.43 [95% CI: 28.54-30.32]; 8 larvae: 27.85 [95% CI: 26.0-29.70]; 16 larvae: 26.53 [95% CI: 24.93-28.12]; 32 larvae: 25.31 [95% CI: 23.32-27.30] (Figure 3B).

**Figure 3.**
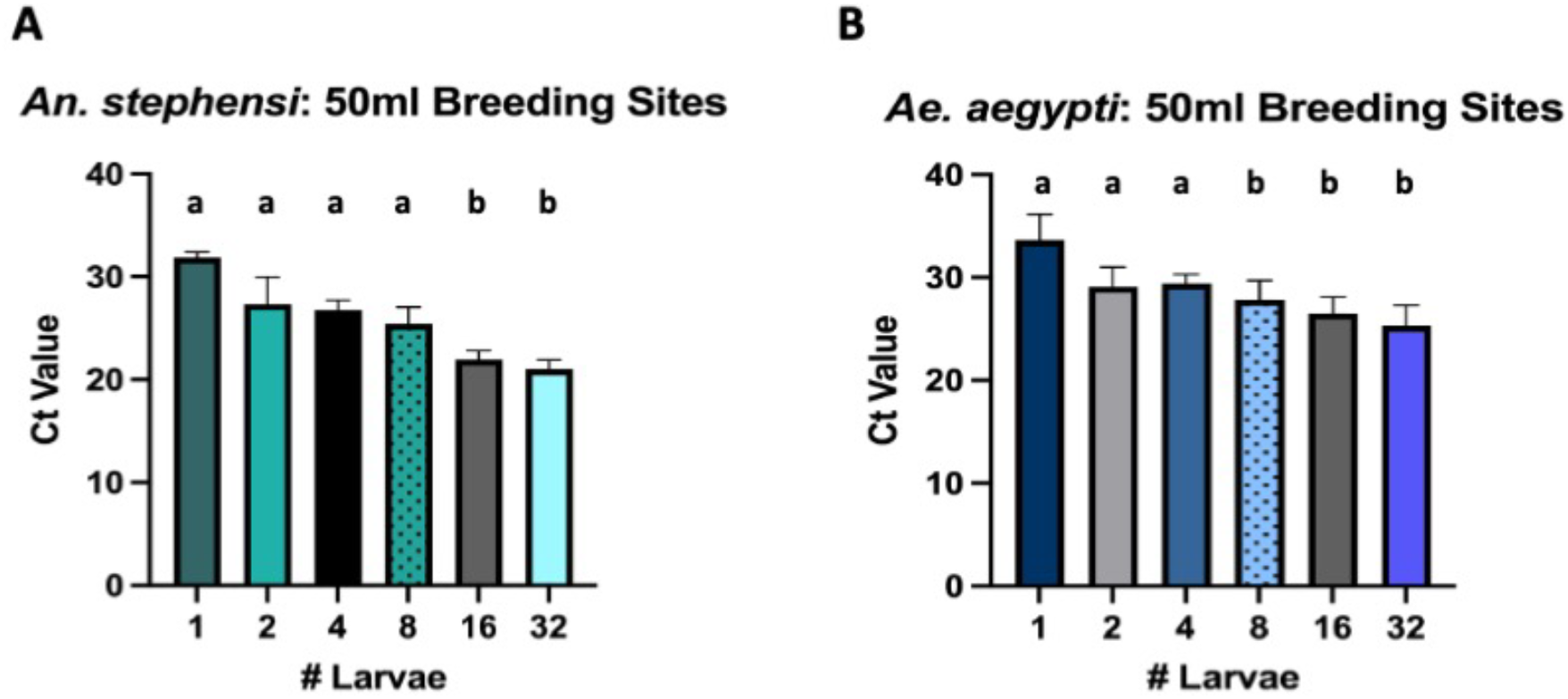
Taqman qPCR detection of *An. stephensi* (A) and *Ae. aegypti* (B) from 50ml breeding sites. qPCR detection for all extractions were run in technical triplicate. Conditions sharing a superscript do not differ significantly (Dunn’s multiple comparisons test, *p*>0.05). Error bars indicate 95% confidence intervals (CIs).

Similarly, a dose response was detectable for both vector species in 1L artificial breeding sites (Figure 2, Experiment 2) with increasing larval densities; unsurprisingly, by comparison to the 50ml breeding sites, Ct values were generally higher at each larval density (Figure 4A and B). The average Ct values for *An. stephensi* for each larval density were 1 larva: 37.82 [95% CI: 36.16-39.47]; 2 larvae: 34.90 [95% CI: 33.27-36.53]; 4 larvae: 33.06 [95% CI: 32.14-33.98]; 8 larvae: 25.32 [95% CI: 23.44-27.21]; 16 larvae: 21.71 [95% CI: 20.08-23.33]; 32 larvae: 21.86 [95% CI: 20.95-22.78] (Figure 4A). The average Ct values for *Ae. aegypti* for each larval density were 1 larva: 37.70 [95% CI: 36.72-38.68]; 2 larvae: 34.91 [95% CI: 33.18-36.65]; 4 larvae: 32.68 [95% CI: 32.20-33.16]; 8 larvae: 30.57 [95% CI: 30.26-30.89]; 16 larvae: 28.80 [95% CI: 28.47-29.12]; 32 larvae: 28.54 [95% CI: 27.58-29.50] (Figure 4B).

**Figure 4.**
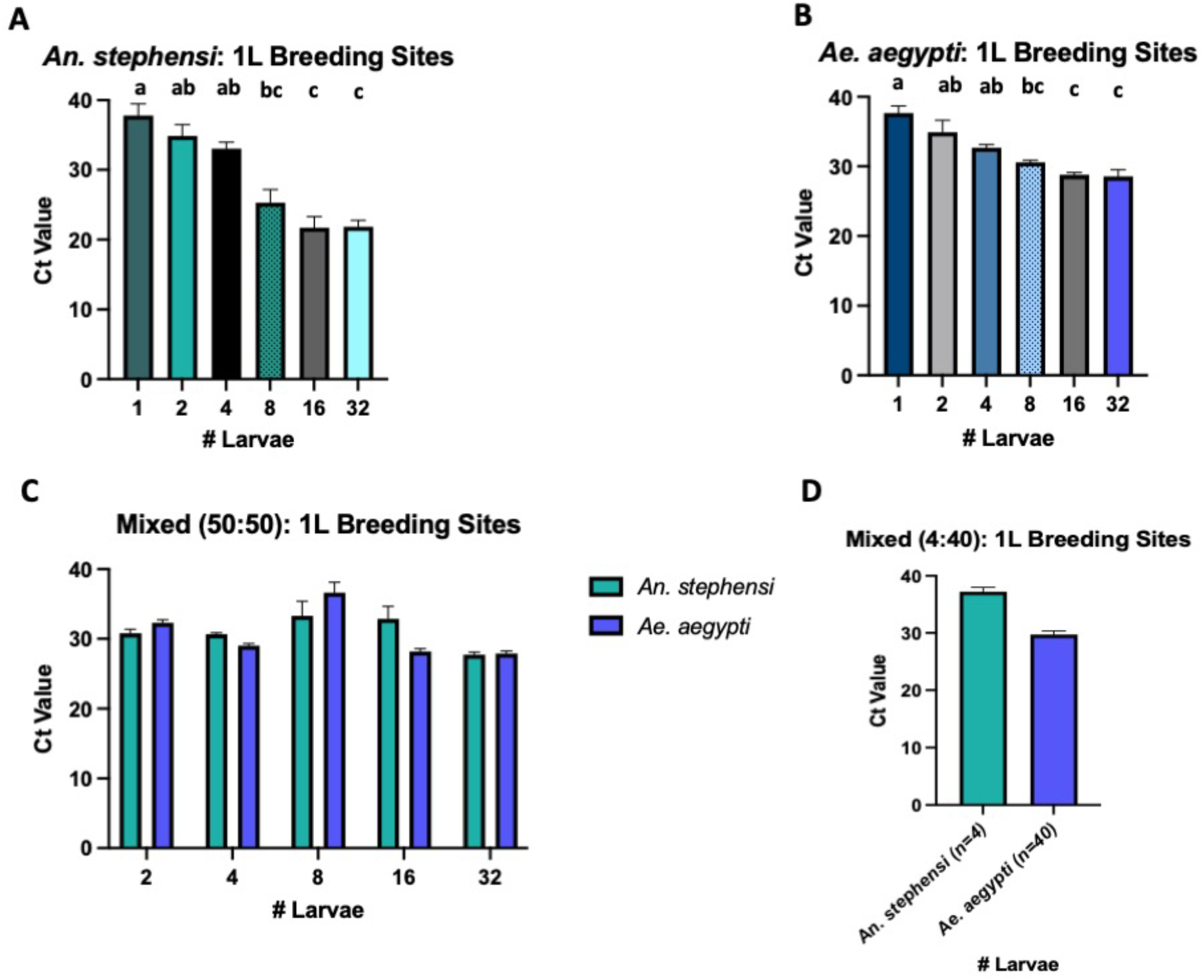
Taqman qPCR detection of *An. stephensi* (A), *Ae. aegypti* (B) or both species in equal (C) or 4:40 (D) proportions from 1L breeding sites. qPCR detection for all extractions were run in technical triplicate. Conditions sharing a superscript do not differ significantly (Dunn’s multiple comparisons test, *p*>0.05). Error bars indicate 95% confidence intervals (CIs).

In 1L artificial breeding sites with mixed vector species in equal proportions (Figure 2, Experiment 2), Ct values were generally lower than those detected in 1L breeding sites containing individual species at lower larval densities (Figure 4C). The average Ct values for *An. stephensi* in mixed breeding sites at each larval density were: 2 larvae: 30.79 [95% CI: 30.22-31.35]; 4 larvae: 30.65 [95% CI: 30.44-30.86]; 8 larvae: 31.57 [95% CI: 31.17-31.96]; 16 larvae: 31.84 [95% CI: 29.53-34.15]; 32 larvae: 27.71 [95% CI: 27.35-28.07] (Figure 4C). The average Ct values for *Ae. aegypti* in mixed breeding sites at each larval density were: 2 larvae: 32.29 [95% CI: 31.88-32.71]; 4 larvae: 29.01 [95% CI: 28.71-29.31]; 8 larvae: 36.63 [95% CI: 35.16-38.10]; 16 larvae: 28.18 [95% CI: 27.78-28.58]; 32 larvae: 27.88 [95% CI: 27.51-28.26] (Figure 4C).

In 1L artificial breeding sites with mixed vector species at a 4:40 ratio (*An. stephensi:Ae. aegypti*), both species were still detectable, with higher average Ct values for the minority species (*An. stephensi*); the average Ct value for *An. stephensi* was 37.20 [95% CI: 36.43-37.97] and for *Ae. aegypti* was 29.72 [95% CI: 29.08-30.36] (Figure 4D).

### Rate of An. stephensi eDNA degradation in artificial breeding sites

A longitudinal cohort of 1L artificial breeding sites were monitored for 2 weeks, following the emergence of 32 *An. stephensi* pupae, to observe the rate of eDNA degradation. A negative dose response was detectable with increasing Ct values over time (Figure 5).The average Ct values for *An. stephensi* eDNA at each time point were 24 hours: 28.90 [95% CI: 27.47-30.32]; 48 hours: 28.31 [95% CI: 27.99-28.63]; 72 hours: 28.97 [95% CI: 28.53-29.41]; 96 hours: 29.72 [95% CI: 29.45-29.99]; 120 hours: 29.48 [95% CI: 28.92-30.04]; 144 hours: 31.19 [95% CI: 30.49-31.88]; 168 hours: 31.38 [95% CI: 31.05-31.71]; 192 hours: 31.56 [95% CI: 31.17-31.94]; 216 hours: 32.82 [95% CI: 32.30-33.34]; 240 hours: 32.57 [95% CI: 31.93-33.20]; 288 hours: 36.96 [95% CI: 36.43-37.50] (Figure 5). *An. stephensi* eDNA was undetectable by qPCR after 312 hours (13 days).

**Figure 5.**
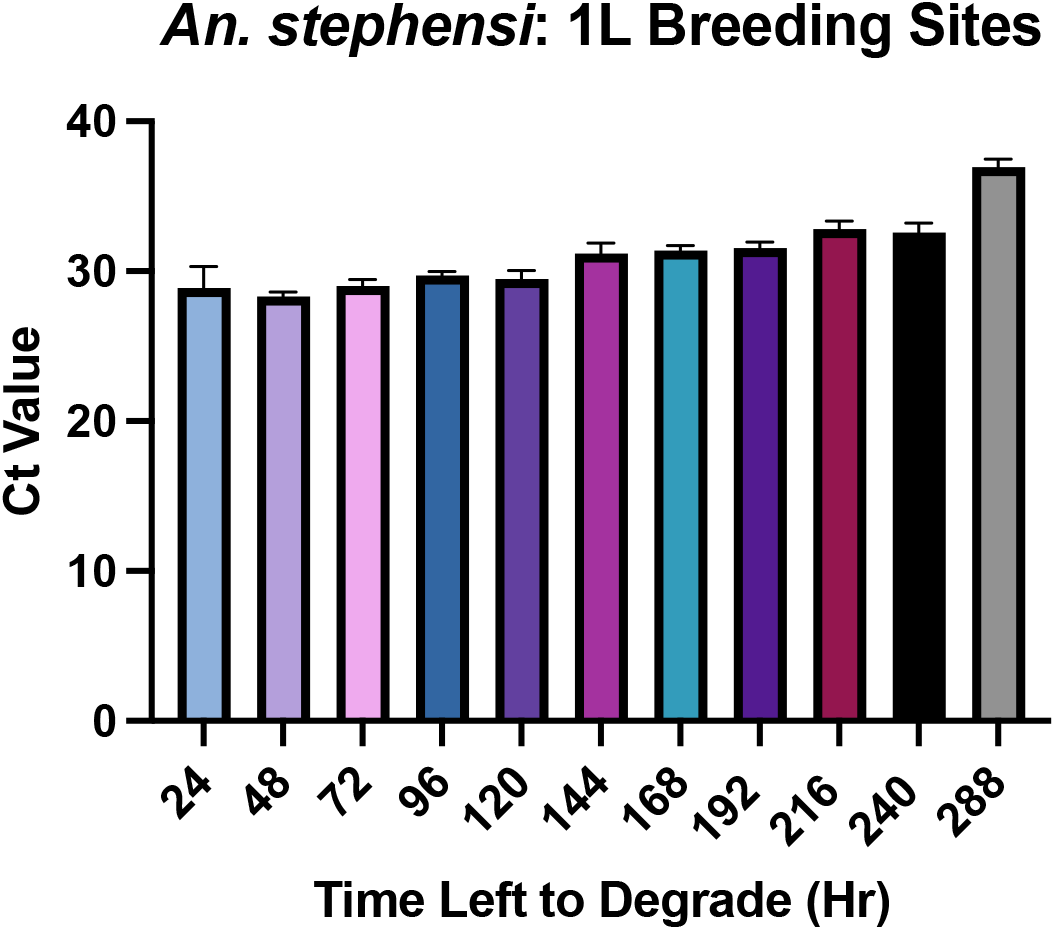
Rate of *An. stephensi* eDNA degradation over 14 days. qPCR detection for all extractions were run in technical triplicate. Error bars indicate 95% confidence intervals (CIs).

### Detection of insecticide resistance genes from eDNA in artificial breeding sites

Characterisation of published insecticide resistance mechanisms in *An. stephensi* using amplicon-sequencing, was undertaken with eDNA extracted from 50ml artificial breeding sites. Amplicons covering the *voltage-gated sodium channel* (*vgsc*), *glutathione-s-transferase 2* (*GSTe2*) and *acetylcholinesterase-1* (*ace-I*) genes, which contain genetic variants associated with insecticide resistance, were tested (Acford-Palmer *et al*. in prep.).

Of the 6 eDNA samples tested (containing 1, 2, 4, 8, 16, and 32 larvae), no PCR amplicons were generated for either eDNA-L1 or -L2 samples. Average read depth per amplicon for eDNA-L4, -L8, -L16 and -L32 is presented in Table 1; eDNA-L32 was the only sample that produced all 7 amplicons when imaged on an agarose gel. The Ace-1 II amplicon for eDNA-L4 was the only incidence of a sample which produced an average coverage too low for SNP calling (>50). Primer pairs for VGSC II, III and IV amplicons proved the least sensitive, with only eDNA-L32 yielding positive results (Table 1).

**Table 1.** Coverage for *An. stephensi* eDNA-L4, -L8, -L16, and -L32 amplicon targets. Chromosome number (chr), start and end of the target amplicon, the target gene, amplicon name and coverage are displayed.

| Chr | Start | End | Target Gene | Amplicon | L4 | L8 | L16 | L32 |
| --- | --- | --- | --- | --- | --- | --- | --- | --- |
| 2 | 60913833 | 60914331 | *ace-1* | ACE-1_I | 4.12 | 43.42 | 67.90 | 836.42 |
| 2 | 60913236 | 60913796 | *ace-1* | ACE-1_II | 7021.93 | 3620.82 | 3163.21 | 2547.49 |
| 3 | 42830105 | 42830628 | *vgsc* | VGSCI | 58.99 | 253.52 | 12.91 | 1032.54 |
| 3 | 42817282 | 42817823 | *vgsc* | VGSCII | 2980.09 | 1117.36 | 1067.52 | 685.12 |
| 3 | 42809619 | 42810112 | *vgsc* | VGSCIII | 0.00 | 0.00 | 0.00 | 140.11 |
| 3 | 42808876 | 42809271 | *vgsc* | VGSCIV | 0.00 | 0.00 | 0.00 | 248.51 |
| 3 | 70580282 | 70580771 | *GSTe2* | GSTe2 | 0.00 | 0.00 | 0.00 | 281.57 |

A total of 40 variants were identified accounting for 33 single nucleotide polymorphisms (SNPs) and 7 insertions or deletions (indels). Only one missense mutation was identified in the *Ace-1* gene in amino acid 177; this SNP was present in the eDNA-L8 and eDNA-L16 samples, with eDNA-L8 genotyped as heterozygous and eDNA-L16 as the homozygous alternate. This SNP has been previously reported in both colony and wild-caught samples (Acford-Palmer *et al*. in prep.). No SNPs associated with insecticide resistance were identified.

Similarly, eDNA from *Ae. aegypti* 50ml artificial breeding sites containing 4 and 8 larvae did not amplify using amplicon-seq primers, targeting the *vgsc* and *ace-1* genes (Collins *et al*. in prep.). PCR reactions using eDNA from *Ae. aegypti* artificial breeding sites containing 16 larvae (eDNA-L16) were able to amplify 5 targets: *ace-1* and four amplicons for the *vgsc* (DomainIVS6, DomainIV, Domain IIIExon36 and DomainIIIExon35; Table 2).

**Table 2.** Coverage for *Ae. aegypti* eDNA-L16 insecticide resistant amplicon targets. Chromosome number (chr) start and end of the target amplicon, the target gene, amplicon names and their respective coverage are displayed.

| Chr | Start | End | Target Gene | Amplicon | eDNA-L16 coverage |
| --- | --- | --- | --- | --- | --- |
| 3 | 161500008.00 | 161500475.00 | *ace1* | Ace1 | 24756.60 |
| 3 | 315931391.00 | 315931846.00 | *vgsc* | DomainIVS6 | 32260.40 |
| 3 | 315931739.00 | 315932233.00 | *vgsc* | DomainIV | 6567.40 |
| 3 | 315938703.00 | 315939191.00 | *vgsc* | DomainIIIExon36 | 1391.40 |
| 3 | 315939789.00 | 315939789.00 | *vgsc* | DomainIIIExon33_34 | 0.00 |
| 3 | 315983627.00 | 315984124.00 | *vgsc* | DomainII | 0.00 |
| 3 | 315983774.00 | 315984267.00 | vgsc | DomainIIExon26 | 0.00 |
| 3 | 315998366.00 | 315998809.00 | vgsc | DomainIIS4 | 0.00 |
| 3 | 316080615.00 | 316081078.00 | vgsc | DomainI | 6.30 |
| 3 | 315938919.00 | 315939398.00 | vgsc | DomainIIIExon35 | 790.40 |

Analysis of the DNA sequences obtained for the eDNA-L16 sample did not show any mutations. The sample coverage for each target amplicon is displayed in Table 2 and showed very good coverage ranging from 790.4-to 32260.40-fold. Five targets, however, did not amplify or showed coverage below the cut-off of 30 (DomainIIIExon33_34, DomainII, DomainIIExon26, DomainIIS4 and DomainI). Differences in gene amplification may reflect differences in relative primer efficiency.

## Discussion

The invasion and establishment of *An. stephensi* in the Horn of Africa represents an imminent and significant regional threat, which may jeopardise malaria control, particularly in urban areas which were formally free from disease transmission^24,25^. To develop novel methods of vector surveillance, this study evaluated the feasibility of using eDNA for simultaneous detection of *An. stephensi* and *Ae. aegypti*, which have both been observed co-habiting in natural breeding sites in Ethiopia^23^. Study findings demonstrated that *An. stephensi* and *Ae. aegypti* eDNA deposited by as few as one second instar larvae in 1L of water was detectable by qPCR. In general, dose responses were evident for detection levels of both vector species, with increasing densities of larvae, resulting in significantly lower Ct values, with some degree of stochastic variation observed. This aligns with previous laboratory studies which demonstrated similar levels of detection of *An. gambiae* sensu stricto in artificial breeding sites^30^. Furthermore, the multiplex TaqMan assay developed in this study displayed comparable levels of detection between both vector species, when used to amplify eDNA from mixed breeding sites, in a 50:50 or 1:10 *An. stephensi:Ae. aegypti* ratio in 1L of water.

*Anopheles* breeding sites can be highly heterogenous in both their ecology and temporality, which will directly affect the sensitivity of eDNA detection. In rural environments, breeding sites commonly comprise rice paddies, ponds, puddles, village pumps and associated troughs, cesspools, water basins, agricultural trenches, dams, edges of rivers/streams, in animal/human footprints and irrigation canals and drains^32–34^. In urban landscapes any natural or man-made feature that collects any form of stagnant water is a potential breeding site, particularly in and around deteriorating infrastructure, such as broken water pipes, blocked drains, construction sites, lorry tyre tracks and catch pits^35^. *An. gambiae* s.l. breeding sites are frequently freshwater habitats that are small, temporary, clean and sun-exposed, although some studies suggest that these larvae can survive in dirty and polluted environments^36,37^; while *An. funestus* s.l. tend to breed in larger semi-permanent water bodies containing aquatic vegetation and algae^38^. In Ethiopia, *An. stephensi* has been found in both rural and urban areas, breeding in *Aedes*-type sites (e.g. artificial water containers, buckets and tanks)^23^. With such variation in breeding site environments, it is anticipated that the rate of eDNA persistence will differ quite considerably, with respect to water turbidity, velocity, UV exposure, salinity/pH, pollution, and co-occupancy (e.g. with other insects, mosquito vectors, fish or tadpoles). In our study, *An. stephensi* eDNA, derived from emergent pupae for 24 hours, was remarkably stable, and still detectable almost two weeks later. Similarly, eDNA from 15 *Ae. albopictus* larvae, reared for 12 days in 500ml water, was demonstrable up to 25 days after live material was removed^31^. Our study may have in fact underestimated the sensitivity of eDNA detection, by only allowing larvae to remain in artificial breeding sites for 24 hours, as well as the rate of eDNA degradation, due to highly controlled insectary conditions (i.e. constant temperature, humidity and light:dark cycles).

In addition to vector species identification, this study investigated the potential to exploit eDNA for molecular insecticide resistance surveillance. Using recently developed amplicon-seq panels for *An. stephensi* (Acford-Palmer *et al*. in prep.) and *Ae. aegypti* (Collins *et al*. in prep.), fragments of key insecticide resistance genes (portions of the voltage-gated sodium channel and acetylcholinesterase) were able to be amplified from as few as 16-32 second instar larvae in 50ml of water; as expected, no insecticide resistance mutations were identified in our insecticide-susceptible insectary colonies. The detection sensitivity of insecticide resistance genes in natural breeding sites may be greater than in our laboratory study, particularly as larval densities can be higher and vector populations will deposit eDNA throughout their ~2 week lifecycle in breeding sites. Conventional insecticide resistance monitoring is based on larval dipping from known, productive breeding sites, rearing of vectors to the adult stage, followed by phenotypic characterization in bioassays and/or genotypic analysis using various molecular techniques^39^. eDNA detection offers a new avenue to explore molecular insecticide resistance at the community-level, circumventing some of the laborious sampling requirements and potentially allow wider-scale surveillance of water bodies which may be less productive sites or entirely unknown to local entomologists. This strategy could be combined with additional analytical techniques, such as high-performance liquid chromatography, to survey the extent of breeding site contamination with agricultural pesticides, a known driver of insecticide resistance selection in *Anopheles* populations^40,41^.

Water sampling for mosquito eDNA detection represents a cheap, low-technology tool, which requires virtually no entomological skills or training, and thus has the potential to be easily integrated into citizen science initiatives. Furthermore, with regards to *An. stephensi* specifically, eDNA detection could be considered for surveillance of seaports in countries at greatest risk of introduction of this invasive species^42^. Following laboratory validation in this study, further work is required to assess the feasibility of sampling eDNA of *An. stephensi/Ae. aegypti in situ*, including an assessment of environmental risk factors which impact eDNA detection levels and comparison of eDNA quantification *versus* larval density per breeding site.

## Conclusions

This study validated the use of eDNA for simultaneous detection of *An. stephensi* and *Ae. aegypti* in shared artificial breeding sites. Study findings demonstrated that *An. stephensi* and *Ae. aegypti* eDNA deposited by as few as 1 second instar larva in 1 litre of water was detectable. Characterization of molecular insecticide resistance mechanisms, using novel amplicon-seq panels for both vector species, was possible from eDNA derived from as few as 16-32 second instar larvae in 50ml of water. eDNA surveillance has the potential to be implemented in local endemic communities as part of citizen science initiatives, and/or in cargo ports at points of country entry, to monitor the spread of invasive malaria vector species. Further studies are required to validate the feasibility of this technique under field conditions.

## Materials and Methods

Laboratory-reared second instar *An. stephensi* (SD500 strain) and *Ae. aegypti* (AEAE strain) larvae were used in all experiments. The larvae were reared in sterile, distilled water, in plastic bowls (20 × 18 × 7 cm) under controlled insectary conditions (26–28°C, relative humidity 70– 80 % and 12:12 hour light:dark cycles) and fed once daily with NISHIKOI staple fish food pellets (Nishikoi, UK).

### Experiment 1: 50ml artificial breeding sites

In the first experiment, 18 different conditions were tested by adding 50ml of sterile, distilled, water to 18 sterile 50ml falcon tubes (Fisher Scientific, UK). We performed three biological replicates with 6 different larval densities: 1, 2, 4, 8, 16 and 32 of either *An. stephensi* or *Ae. aegypti* larvae (Figure 1). Three negative control experimental habitats with no larvae were run in parallel for each condition. Larvae were left in each 50ml breeding site for 24 hours, prior to manual removal, using a sterile pipette.

### Experiment 2: 1L artificial breeding sites

In the second experiment, 18 different conditions were tested by adding 1L of sterile, distilled water to 18 sterile plastic rectangular containers. We performed three biological replicates with 6 different larvae densities: 1, 2, 4, 8, 16 and 32 larvae. Three negative control experimental habitats with no larvae were run in parallel for each condition. Larvae were left in each 1L breeding site for 24 hours, prior to manual removal, using a sterile pipette. *An. stephensi* and *Ae. aegypti* were evaluated initially in separate 1L breeding sites and then in mixed breeding sites. The mixed breeding site experiments used 18 different conditions, comprising three biological replicates with 5 larvae densities: 2, 4, 8, 16 and 32 larvae of *An. stephensi* and *Ae. aegypti* in equal proportions (50:50) and three biological replicates with larvae of the two species in different proportions: 4:40 *An. stephensi:Ae. aegypti* larvae.

### Experiment 3: eDNA degradation in artificial breeding sites

In the third experiment, the rate of eDNA degradation in artificial breeding sites was assessed. One litre of sterile, distilled water was added to 9 sterile plastic rectangular containers. We performed three biological replicates with 32 *An. stephensi* pupae added to each container. Three negative control experimental habitats with no pupae were run in parallel for each condition. Pupae were left in each 1L breeding site for 24 hours, prior to manual removal of all emerged adults, remaining pupae and pupal skins, using a sterile pipette. Water samples were then removed daily for 14 days, following the removal of adults and pupae.

### eDNA Extraction

eDNA from water samples were concentrated prior to extraction. For each breeding site replicate, 15ml of water was added to a sterile 50ml falcon tube (Fisher Scientific, UK) and 1.5ml of 3M sodium acetate solution (pH 5.2) (Sigma-Aldrich, UK) was immediately added, followed by 33ml of absolute ethanol and stored overnight at −20°C. Samples were then centrifuged at 8000 rpm at 6°C for 30 minutes. The supernatant was discarded, and the pellet was washed in 20ml of absolute ethanol by centrifuging at 8000 rpm at 6°C for 10 minutes. The supernatant was discarded, and ethanol allowed to evaporate at 65°C. The pellet was dissolved in 720μl ATL buffer and 80μl proteinase K (Qiagen, UK) and incubated overnight at 56°C. eDNA was extracted using a Qiagen DNeasy 96 Blood and Tissue kit (Qiagen, UK), according to the manufacturer’s protocol, with minor modifications.

### An. stephensi and Ae. aegypti eDNA PCR primer design and validation

We designed a multiplex TaqMan assay to distinguish between *An. stephensi* (ASTE016222) and *Ae. aegypti* (AAEL018662) eDNA based on a fragment of cytochrome oxidase I (COX1). Forward (5’-CAGGAATTACWTTAGACCGAMTACC-3’) and reverse (5’-TCAAAATAARTGTTGRTATAAAATRGGGTC-3’) primers and two TaqMan fluorescence-labelled probes (*An. stephensi* probe 5’-/5HEX/ATTACTATA/ZEN/TTACTTACAGACCG/3IABkFQ/-3’ and *Ae. aegypti* probe 5’-/56FAM/ATTACTATG/ZEN/TTATTAACAGACCG/3IABkFQ/-3’) were designed using Geneious Prime® 2021.1.1 to amplify a 202 bp region of COXI (Figure 4). Primers and probes were selected by reference to all available COXI sequences from NCBI GenBank (n=4402) for *Ae. aegypti, An. arabiensis, An. gambiae sensu stricto, An. pharoensis, An. quadriannulatus, An. funestus, An. stephensi* and *Cx. quinquefasciatus*, which were used to confirm the presence/absence of species-specific SNPs and to account for additional intra-species genetic diversity.

Standard curves of Ct values for each probe were generated individually and then multiplexed using a ten-fold serial dilution of control *An. stephensi* or *Ae. aegypti* DNA (extracted from SD500 and AEAE laboratory colonies, respectively) to assess PCR efficiencies. Genomic DNA concentrations were determined using the Qubit 4 fluorometer 1X dsDNA HS assay (Invitrogen, UK). Standard curve reactions were performed in a final volume of 10μl containing 2X PrimeTime® Gene Expression Master Mix (IDT, USA), 250nM of forward and reverse primers and 150nM of each probe and 2μl genomic DNA. Reactions were run on a Stratagene Mx3005P Real-Time PCR system (Agilent Technologies) at 95°C for 3 minutes, followed by 40 cycles of 95°C for 15 seconds and 60°C for 1 minute. All assays were run in technical triplicate alongside PCR no-template controls (NTCs).

To confirm primer and probe specificity, the multiplex assay was used to assess amplification of stock genomic DNA from other medically important sympatric vector species: *An. stephensi, Ae. aegypti, An. arabiensis, An. gambiae* s.s., *An. funestus* and *Cx. quinquefasciatus*.

### Detection of An. stephensi and Ae. aegypti eDNA from artificial breeding sites

Following eDNA extraction, eDNA detection was performed in a final volume of 10μl containing 2X PrimeTime® Gene Expression Master Mix (IDT, USA), 250nM of forward and reverse primers, 150nM of each probe and 4.2μl eDNA. Reactions were run on a Stratagene Mx3005P Real-Time PCR system (Agilent Technologies) at 95°C for 3 minutes, followed by 40 cycles of 95°C for 15 seconds and 60°C for 1 minute. All assays were run in technical triplicate alongside PCR no-template controls (NTCs).

### qPCR data analysis

Stratagene MxPro qPCR software (Agilent Technologies, UK) was used to produce qPCR standard curves. qPCR assay limits of detection (LoD) were determined using the “Generic qPCR LoD / LoQ calculator”^43^, implemented in R version 4.0.2^44^. All other statistical analyses were conducted in GraphPad Prism v9.4.0.

### Amplicon primer design and PCR amplification

To characterize molecular mechanisms of insecticide resistance in *An. stephensi*, briefly, three genes of interest were identified, and a total of 7 amplicon primer pairs were designed to target SNPs, previously associated with insecticide resistance (Acford-Palmer *et al*. in prep.). To enable sample multiplexing, each primer had a 6bp barcode attached to the 5’ end to allow for sample identification in downstream analysis. Amplicons were generated using the eDNA extracted from experiment #1 (1, 2, 4, 8, 16, and 32 larvae in 50ml artificial breeding sites). PCR reactions contained 5X Q5 reaction buffer (New England Biolabs, UK), Q5® High-Fidelity DNA polymerase (New England Biolabs, UK), 250nM of forward and reverse primers and 2μl of eDNA. PCR reaction conditions were an initial 30 second denaturation at 95°C, followed by 35 cycles of 98°C for 10 seconds, 60°C for 30 seconds and 72°C for 30 seconds, and a final elongation step at 72°C for 2 minutes. PCR products were visualized on SYBR Safe 1% agarose gels (Invitrogen, UK), prior to purification using Agencourt AMPure XP magnetic beads (Beckman Coulter, UK), using a ratio of 0.7:1 (μl of beads to DNA).

To characterize molecular mechanisms of insecticide resistance in *Ae. aegypti*, briefly, sequences for portions of the voltage-gated sodium channel (*vgsc*; AAEL023266-RL), acetylcholinesterase-1 and (*Ace-1*; AAEL000511-RJ) were extracted from publicly available assemblies for *Ae. aegypti* (LVP AGWG) (Collins *et al*. in prep.). Forward and reverse primers were designed using PrimerBLAST software to amplify regions of 450-500 bp that contain known SNP loci or regions of interest, with 6bp inline barcodes and partial Illumina tails that allow the samples to be sequenced on an Illumina sequencing platform. PCR reactions contained 5X Q5 reaction buffer (New England Biolabs, UK), Q5® High-Fidelity DNA polymerase (New England Biolabs, UK), 250nM of forward and reverse primers and 2μl of eDNA extracted from experiment #1 (1, 2, 4, 8 and 16 larvae in 50ml artificial breeding sites). PCR reaction conditions were 98°C for 30 seconds, followed by 35 cycles of 98°C for 10 seconds, 57 °C for 60 seconds and 72°C for 90 seconds. PCR products were visualized on SYBR Safe 1% agarose gels (Invitrogen, UK), prior to purification using Agencourt AMPure XP magnetic beads (Beckman Coulter, UK), using a ratio of 0.7:1 (μl of beads to DNA).

### Amplicon sequencing

The concentration of purified PCR products was measured using the Qubit 4 fluorometer 1X dsDNA HS assay (Invitrogen, UK) and samples with unique barcode combinations were pooled in equal concentrations to create an overall pool of 20ng/μl in 25μl total volume. A second PCR to insert Illumina adaptors (no further library preparation required) and amplicon sequencing was performed at Genewiz (Illumina-based Amplicon-EZ service).

### Amplicon sequencing analysis

Samples were demultiplexed based on the in-line barcodes in each forward and reverse primer using an in-house python script (Collins *et al*. in prep.). Sequences were trimmed, aligned to reference (*Ae. aegypti* LVP AGWG for *Ae. aegypti* or UCI_ANSTEP_V1 assembly for *An. stephensi*) and data was quality checked using FastQC and Samclip. Paired-end reads were mapped against the reference sequences using the BWA-MEM algorithm. Variants were then called using three packages: Freebayes, GATK, and an in-house naive variant caller. Identified variants were then compared and normalised against each other to reduce false negatives and positives, and then filtered and annotated using BCFtools; for *An. stephensi*, variants were annotated using snpEFF, based on a manually built database for the UCI *An. stephensi* genome assembly. SNPs were then manually filtered to ensure that they appeared in more than one sample, with an allele depth of >50.

**Figure 4.**
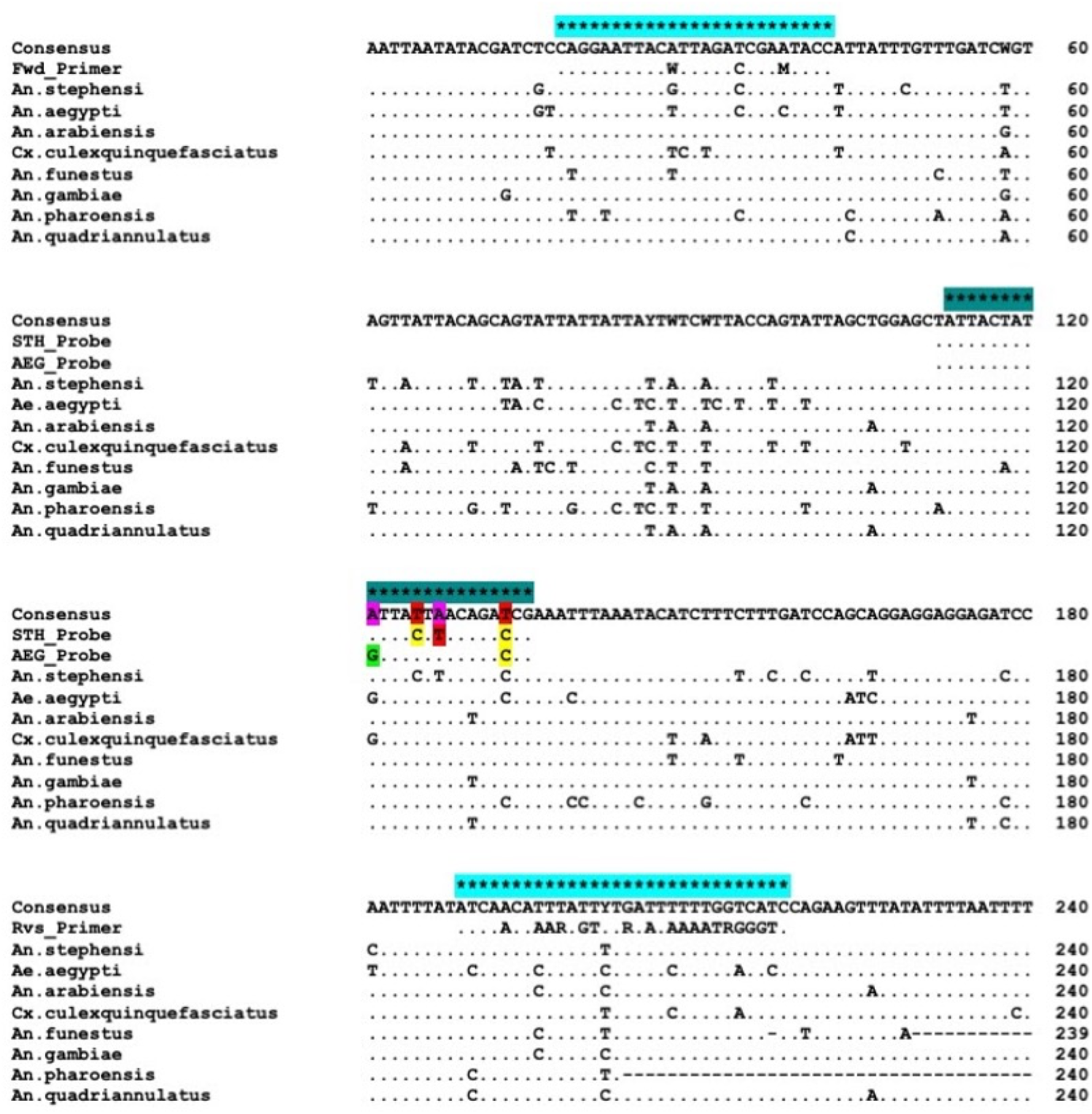
Consensus alignment of reference COXI sequences used for primer and probe design. Positions of forward and reverse primers are highlighted by light blue asterisks. Species-specific probes are highlighted in teal, with single-nucleotide polymorphisms in colour.

## Data availability

The qPCR datasets generated during this study are contained within the supplementary material. Sequence data generated by this study will be available at Sequence Read Archive (SRA) BioProject upon publication.

## Authors’ contributions

MK, SC, and LAM designed the study. MK led the entomological experiments with support from NMP and BP. LAM led the qPCR analysis. HAP, MOC, EC and SC performed the amplicon sequencing. TW provided laboratory resources. JP and TGC performed the bioinformatics analysis. SC and JL provided project oversight and funding. MK, HAP, MOC and LAM drafted the manuscript, which was revised by co-authors. All authors read and approved the final manuscript.

## Acknowledgements

Study funding was provided by the UK Foreign, Commonwealth and Development Office [Health Research Programme Consortia (RPCs): RAFT (Resilience Against Future Threats through Vector Control), PO8615] and a Wellcome Trust/Royal Society Sir Henry Dale Fellowship awarded to TW (101285/Z/13/Z): https://wellcome.org and https://royalsociety.org.

## References

1 Global vector control response 2017-2030. Global vector control response 51 (2017).

2 Organization, W. H. World Malaria Report 2021. (World Health Organization, Geneva, 2021).

3 Collaborators, L. B. o. D. N. T. D. The global distribution of lymphatic filariasis, 2000-18: a geospatial analysis. Lancet Global Health 8, e1186–e1194 (2020).

4 Mordecai, E. A., Ryan, S. J., Caldwell, J. M., Shah, M. M. & LaBeaud, A. D. Climate change could shift disease burden from malaria to arboviruses in Africa. Lancet Planetary Health 4, e416–e423 (2020).

5 Simo, F. B. N. et al. Dengue virus infection in people residing in Africa: a systematic review and meta-analysis of prevalence studies. Scientific Reports 9, 13626 (2019).

6 Buchwald, A. G., Hayden, M. H., Dadzie, S. K., Paull, S. H. & Carlton, E. J. Aedes-borne disease outbreaks in West Africa: a call for enhanced surveillance. Acta Tropica 209, 105468 (2020).

7 Ushijima, Y. et al. Surveillance of the major pathogenic arboviruses of public health concern in Gabon, Central Africa: increased risk of West Nile virus and dengue virus infections. BMC Infectious Diseases 21, 265 (2021).

8 Bowman, L. & McCall, P. J. Evaluating the evidence for effectiveness of vector control of dengue outbreaks by systematic review and meta-analysis. American Journal of Tropical Medicine and Hygiene 91, 231 (2014).

9 Organization, W. H. Malaria surveillance, monitoring and evaluation: a reference manual. (World Health Organization, Geneva, 2018).

10 Runge-Ranzinger, S., McCall, P. J., Kroeger, A. & Horstick, O. Dengue disease surveillance: an updated systematic literature review Tropical Medicine & International Health 19, 1116–1160 (2014).

11 Cansado-Utrilla, C. et al. An assessment of adult mosquito collection techniques for studying species abundance and diversity in Maferinyah, Guinea. Parasites & Vectors 13, 150 (2020).

12 Abong’o, B. et al. Comparison of four outdoor mosquito trapping methods as potential replacements for human landing catches in western Kenya. Parasites & Vectors 14, 320 (2021).

13 Programme, W. H. O. G. M. Vector alert: Anopheles stephensi invasion and spread. (Geneva, 2019).

14 Faulde, M. K., Rueda, L. M. & Khaireh, B. A. First record of the Asian malaria vector Anopheles stephensi and its possible role in the resurgence of malaria in Djibouti, Horn of Africa. Acta Tropica 139, 39–43 (2014).

15 Carter, T. E. et al. First detection of Anopheles stephensi Liston, 1901 (Diptera: culicidae) in Ethiopia using molecular and morphological approaches. Acta Tropica 188, 180–186 (2018).

16 Tadesse, F. G. et al. Anopheles stephensi Mosquitoes as Vectors of Plasmodium vivax and falciparum, Horn of Africa, 2019. Emerging Infectious Diseases 27, 603–607 (2021).

17 Ahmed, A. et al. Invasive Malaria Vector Anopheles stephensi Mosquitoes in Sudan, 2016-2018. Emerging Infectious Diseases 27, 2952–2954 (2021).

18 Ahmed, A., Khogali, R., Elnour, M. B., Nakao, R. & Salim, B. Emergence of the invasive malaria vector Anopheles stephensi in Khartoum State, Central Sudan. Parasites & Vectors 14, 511 (2021).

19 Organization, W. H. WHO malaria threats map, <https://apps.who.int/malaria/maps/threats/> (

20 Seyfarth, M., Khaireh, B. A., Abdi, A. A., Bouh, S. M. & Faulde, M. K. Five years following first detection of Anopheles stephensi (Diptera: Culicidae) in Djibouti, Horn of Africa: populations established-malaria emerging. Parasitology Research 118, 725–732 (2019).

21 Gutu, M. A. et al. Another dengue fever outbreak in Eastern Ethiopia-An emerging public health threat. PLoS Neglected Tropical Diseases 15, e0008992 (2021).

22 Mulchandani, R. et al. A community-level investigation following a yellow fever virus outbreak in South Omo Zone, South-West Ethiopia. PeerJ 7, e6466 (2019).

23 Balkew, M. et al. Geographical distribution of Anopheles stephensi in eastern Ethiopia. Parasites & Vectors 13, 35 (2020).

24 Sinka, M. E. et al. A new malaria vector in Africa: predicting the expansion range of Anopheles stephensi and identifying the urban populations at risk. Proceedings of the National Academy of Sciences of the United States of America 117, 24900–24908 (2020).

25 Hamlet, A. et al. The potential impact of Anopheles stephensi establishment on the transmission of Plasmodium falciparum in Ethiopia and prospective control measures. BMC Medicine 20, 135 (2022).

26 Organization, W. H. Vector alert: Anopheles stephensi invasion and spread. (World Health Organization, 2019).

27 Thomsen, P. F. & Willerslev, E. Environmental DNA – An emerging tool in conservation for monitoring past and present biodiversity. Biological Conservation 183, 4–18 (2015).

28 Ficetola, G. F., Miaud, C., Pompanon, F. & Taberlet, P. Species detection using environmental DNA from water samples. Biology Letters 4, 423–425 (2008).

29 Harrison, J. B., Sunday, J. M. & Rogers, S. M. Predicting the fate of eDNA in the environment and implications for studying biodiveristy. Proceedingsof the Royal Society B 286, 20191409 (2019).

30 Odero, J., Gomes, B., Fillinger, U. & Weetman, D. Detection and quantification of Anopheles gambiae sensu lato mosquito larvae in experimental aquatic habitats using environmental DNA (eDNA). Wellcome Open Research 3, 26 (2018).

31 Schneider, J. et al. Detection of invasive mosquito vectors using environmental DNA (eDNA) from water samples. PLoS One 11, e0162493 (2016).

32 Zogo, B. et al. Identification and characterization of Anopheles spp. breeding habitats in the Korhogo area in northern Côte d’Ivoire: a study prior to a Bti-based larviciding intervention. Parasites & Vectors 12, 146 (2019).

33 Hamza, A. M. & El Rayah el, A. A Qualitative Evidence of the Breeding Sites of Anopheles arabiensis Patton (Diptera: Culicidae) in and Around Kassala Town, Eastern Sudan. International Journal of Insect Science 8, 65–70 (2016).

34 Ndiaye, A. et al. Mapping the breeding sites of Anopheles gambiae s. l. in areas of residual malaria transmission in central western Senegal. PLoS One 15, e0236607 (2020).

35 Mattah, P. A. et al. Diversity in breeding sites and distribution of Anopheles mosquitoes in selected urban areas of southern Ghana. Parasites & Vectors 10, 25 (2017).

36 Fillinger, U., Sonye, G., Killeen, G. F., Knols, B. G. & Becker, N. The practical importance of permanent and semipermanent habitats for controlling aquatic stages of Anopheles gambiae sensu lato mosquitoes: operational observations from a rural town in western Kenya. Tropical Medicine & International Health 12, 1274–1289 (2004).

37 Walker, K. & Lynch, M. Contributions of Anopheles larval control to malaria suppression in tropical Africa: review of achievements and potential. Medical & Veterinary Entomology 21, 2–21 (2007).

38 Nambunga, I. H. et al. Aquatic habitats of the malaria vector Anopheles funestus in rural south-eastern Tanzania. Malaria Journal 19, 219 (2020).

39 Organization, W. H. Manual for monitoring insecticide resistance in mosquito vectors and selecting appropriate interventions. (Geneva, 2022).

40 Nkya, T. E. et al. Impact of agriculture on the selection of insecticide resistance in the malaria vector Anopheles gambiae: a multigenerational study in controlled conditions. Parasites & Vectors 7, 480 (2014).

41 Hien, A. S. et al. Evidence that agricultural use of pesticides selects pyrethroid resistance within Anopheles gambiae s.l. populations from cotton growing areas in Burkina Faso, West Africa. PLoS One 12, e0173098 (2017).

42 Ahn, J., Sinka, M., Irish, S. & Zohdy, S. Using marine cargo traffic to identify countries in Africa with greatest risk of invasion by Anopheles stephensi. bioRxiv, doi:https://doi.org/10.1101/2021.12.07.471444 (2022).

43 Merkes, C. et al. Reporting the limits of detection (LOD) and quantification (LOQ) for environmental DNA assays. Environmental DNA 2, 271–282, doi:https://doi.org/10.5066/P9AKHU1R. (2020).

44 RStudio: Integrated Development for R. (Bostom, MA, 2020).

